# NOTCH3 active immunotherapy reduces NOTCH3 deposition in brain capillaries in a CADASIL mouse model

**DOI:** 10.1101/2022.07.11.499563

**Authors:** Daniel V. Oliveira, Kirsten G. Coupland, Shaobo Jin, Francesca Del Gaudio, Sailan Wang, Rhys Fox, Julie W. Rutten, Johan Sandin, Johan Lundkvist, Saskia A. J. Lesnik Oberstein, Urban Lendahl, Helena Karlström

**Author notes:** Shared first authorship (Daniel V. Oliveira, Kirsten G. Coupland).

## Abstract

Cerebral autosomal dominant arteriopathy with subcortical infarcts and leukoencephalopathy (CADASIL) is the most common monogenic form of familial small vessel disease and no preventive or curative therapy is available. CADASIL is caused by mutations in the *NOTCH3* gene, resulting in a mutated NOTCH3 receptor, with aggregation of the NOTCH3 extracellular domain (ECD) around vascular smooth muscle cells. In this study we have developed a novel active immunization therapy specifically targeting CADASIL-like aggregated NOTCH3 ECD. Immunizing CADASIL TgN3R182C^150^ mice with aggregates composed of CADASIL-R133C mutated and wild type EGF1-5 repeats for a total of four months resulted in a marked reduction (38-48%) in NOTCH3 deposition around brain capillaries, increased microglia activation and lowered serum levels of NOTCH3 ECD. Active immunization did not impact body weight, general behavior or the number and integrity of vascular smooth muscle cells in the retina, suggesting that the therapy is tolerable. This is the first therapeutic study reporting a successful reduction of CADASIL-like NOTCH3 accumulation in mice supporting further development towards clinical application for the benefit of CADASIL patients.

## Introduction

Cerebral autosomal dominant arteriopathy with subcortical infarcts and leukoencephalopathy (CADASIL, OMIN No. 125310) is the most common monogenic form of cerebral small vessel disease (SVD), affecting approximately 5/100 000 individuals, but is in all likelihood underdiagnosed (Rutten *et al*, 2016). Patients suffering from CADASIL experience migraine with aura, subcortical ischemic events, mood disturbances, apathy, and cognitive impairment (Chabriat *et al*, 2009; Coupland *et al*, 2018; Joutel, 2020; Joutel *et al*, 1996). CADASIL results in white matter lesions, neuronal loss and widespread vascular pathology characterized by degenerating vascular smooth muscle cells (VSMC) and thickening of the arterial wall (fibrosis), which leads to lumen stenosis, for review see (Coupland *et al*., 2018).

CADASIL is exclusively caused by mutations in the *NOTCH3* gene (Joutel *et al*., 1996). The NOTCH3 transmembrane receptor undergoes proteolytic cleavages upon activation by ligands presented on juxtaposed cells, ultimately releasing the Notch intracellular domain (ICD) into the interior of the cell, while the Notch extracellular domain (ECD) is shed from the cell surface (see Fig 1A for details on Notch signaling) (Andersson *et al*, 2011; Coupland *et al*., 2018; Siebel & Lendahl, 2017). The majority of CADASIL mutations are missense mutations confined to the 34 epidermal growth factor (EGF)-like repeats of the NOTCH3 ECD moiety (Rutten *et al*, 2014), resulting in an altered number of cysteine residues in the EGF-like repeats (Fig 1B) (Joutel *et al*., 1996). The CADASIL cysteine-altering mutations perturb the structure of NOTCH3 ECD resulting in NOTCH3 ECD multimerization and aggregation. This in turn leads to the recruitment of microvascular extracellular matrix proteins including metalloproteases and vitronectin, that form the so called granular osmiophilic dense material, GOM, a histopathological hallmark of CADASIL (Capone *et al*, 2016; Duering *et al*, 2011; Karlstrom *et al*, 2002; Monet-Lepretre *et al*, 2013) (Fig 1B). NOTCH3 ECD accumulation is one of the earliest events in CADASIL pathogenesis, indicating that it may cause cellular pathology by inducing changes in the brain microvascular extracellular matrix (Capone *et al*., 2016; Joutel *et al*, 2001; Joutel *et al*, 2010; Monet-Lepretre *et al*., 2013). Collectively, these data argue that CADASIL may be considered an protein misfolding and aggregation disease.

**Figure 1.**
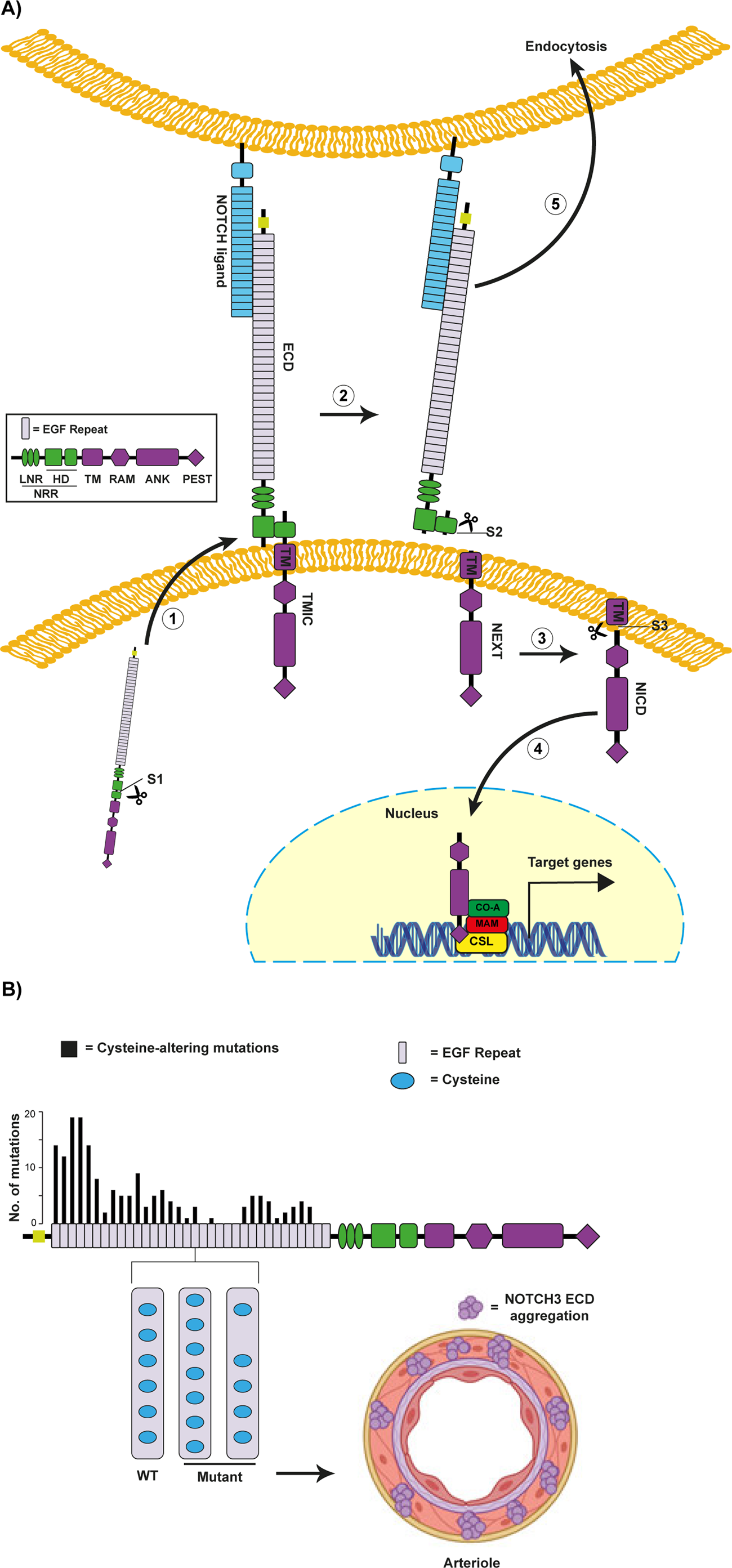
A) Schematic representation of Notch signaling. (1) Furin (S1 cleavage) cleaves the NOTCH3 precursor protein in the Golgi system, resulting in a non-covalently bound heterodimeric protein that is transported to the cell surface. (2) A mechanical traction force is applied to the NOTCH3 ECD when a Notch ligand binds to the EGF repeats 10-11, exposing the extracellular NRR near the cell membrane, which consists of LNR and the heterodimerization domain (in green). Subsequently, ADAM17 cleaves the C-terminal portion of the heterodimerization domain (S2-cleavage). (3) The NEXT, which is made up of a RAM domain, many ANK domains, a PEST domain, and a transmembrane domain, is cleaved by the γ-secretase (S3-cleavage) releasing the N3ICD. (4) The N3ICD binds to the CSL complex protein and together with the co-activator Mastermind-like (MAM) trigger downstream gene transcription in the nucleus. (5) The NOTCH3 ECD and ligand are normally endocytosed by the ligand expressing cell and is degraded in the lysosome. B) Schematic representation of NOTCH3 cerebral autosomal dominant arteriopathy with subcortical infarcts and leukoencephalopathy (CADASIL) mutations. NOTCH3 ECD contains 34 EGF repeat domains, each of which has 6 cysteine residues (WT). Mutations in CADASIL change the number of cysteines to an uneven number of cysteines (Mutant). These unpaired cysteines residues result in incorrect EGF repeat folding, irregular protein folding which leads to an enhanced NOTCH3 ECD multimerization. Distribution of the cysteine-altering mutations that cause CADASIL are shown. In the CADASIL mutant NOTCH3 ECD, the endocytosis is hampered, and NOTCH ECD remains outside of the VSMC and starts to accumulate and aggregate around the vessels. ADAM17, a disintegrin and metalloproteinase domain-containing protein 17; ANK, ankyrin repeats; EGF, epidermal growth factor; HD, heterodimerization domain; LNR, Lin-Notch repeats; PEST, proline (P), glutamic acid (E), serine (S) and threonine (T) degradation domain; RAM, Rbp-associated molecule domain; TM, transmembrane domain.

There are currently no therapies to abrogate or ameliorate the disease process for CADASIL patients but given that NOTCH3 accumulation is a hallmark of the disease, immunotherapy targeting NOTCH3 aggregation or aggregates may be an attractive therapeutic strategy. Immunotherapeutic approaches are gaining increasing attention as novel disease modifying therapies for protein misfolding and aggregation diseases such as certain neurodegenerative diseases (Forman *et al*, 2004; Shrivastava *et al*, 2017). Promising preclinical and clinical results have been obtained in particular for the clearance of Amyloid beta (Aβ) deposits (the so called “senile plaques”) in Alzheimer’s disease (AD). Therapies based on both active and passive immunization can clear amyloid in the brain with great efficiency in preclinical mouse models of Aβ-amyloidosis as well as in patients. Recently, a number of different monoclonal antibodies targeting Aβ aggregates have demonstrated promising clinical improvement in association with amyloid clearance in Phase II and III clinical trials with Aducanumab recently being approved for treatment of AD by the Food and Drug Association (Budd Haeberlein *et al*, 2017; Mintun *et al*, 2021; Tolar *et al*, 2020). Although the efficacy of the compound has been questioned and challenged (Knopman *et al*, 2021), this passive vaccine is still of importance not only for the AD field but also encourages development of immunotherapies for other disorders where protein misfolding and aggregation are believed to play a pivotal pathological role.

CADASIL shares a number of features with AD, including that both are considered protein misfolding and aggregation diseases and that both NOTCH3 and Aβ aggregates accumulate in the extracellular milieu. It is thus conceivable that NOTCH3 aggregates are accessible to efficient antibody-mediated clearance, as has been demonstrated for Aβ amyloid immunotherapies. Thus far, passive immunization has been explored in pre-clinical CADASIL mouse models, with potentially encouraging results (Ghezali *et al*, 2018; Machuca-Parra *et al*, 2017), including ameliorative effects on cerebrovascular dysfunctions such as impaired blood flow and myogenic tone, although no attenuation of NOTCH3 ECD or GOM deposition was observed (Ghezali *et al*., 2018). To directly target the NOTCH3 ECD aggregation, it may therefore be interesting to explore an active immunization strategy, a therapeutic strategy which has been proven safe in several AD trials with different vaccines targeting Aβ (Novak *et al*, 2019; Rosenberg & Lambracht-Washington, 2020; Vandenberghe *et al*, 2017). Active immunization may be particularly appealing as a treatment for CADASIL for a number of reasons. Firstly, the NOTCH3 deposits are located in the vessel walls and thus readily accessible to the circulating humoral immune defense enabling efficient NOTCH3 aggregate targeting. Secondly, given the dominant nature and high penetrance of the NOTCH3 mutations, a large number of CADASIL patients and CADASIL mutation carriers could be identified at a young age, when an active immunization has a greater potential to mount an efficient immune response. Finally, CADASIL is a chronic life-long disease and vaccine injections restricted to a few times yearly, as opposed to monthly or even more frequent injections, which could be the case with passive vaccines, would be advantageous from a patient perspective.

In this study, we report on the development of an active immunization therapy aimed at targeting the CADASIL-associated NOTCH3 pathology as a novel disease-modifying therapy for the treatment of CADASIL. We take advantage of a CADASIL mouse model (TgN3R182C^150^), which expresses a human NOTCH3 R182C receptor and which develops a progressive cerebrovascular NOTCH3 ECD and GOM deposition phenotype in arterioles (Rutten *et al*, 2015), and thus is a suitable preclinical model for the development of therapies targeting CADASIL-associated NOTCH3 pathology. We find that active repeated immunization starting at three months of age results in reduced NOTCH3 ECD deposition around capillaries and lowered levels of NOTCH3 ECD in the blood at seven months of age, which was the end point of the experiments. No loss of VSMC was observed in the retina after NOTCH3 immunization, indicating that the active immunotherapy does not alter normal Notch signaling. Collectively, the data suggest that active immunization that specifically targets aggregated NOTCH3 is an efficacious and tolerable therapeutic strategy for CADASIL therapy development.

## Results

### Production of recombinant N3 EGF_1-5_ antigen for active immunization

In order to perform active immunization in the CADASIL TgN3R182C^150^ mouse model, we first generated a suitable antigen, that would be selective for aggregated NOTCH3 and spare monomeric NOTCH3 receptors, potentially minimizing adverse effects on Notch3 signaling. To this end, we produced an immunogen based on aggregates of the NOTCH3 EGF repeat (EGF1-5), the part of NOTCH3 which contains the majority of all CADASIL-causing mutations identified to date. Although using a mouse model carrying the NOTCH3 R182C mutation, we opted for another cysteine-altering mutation (NOTCH3 R133C) for generation of the aggregated antigen, since this protein previously has been shown to generate aggregates and with a mutation in the “hot-spot” region in the NOTCH3 protein (EGF1-6), but also to determine whether an aggregation-general rather than a mutation-specific therapy could be efficacious. By mixing the NOTCH3 EGF_1-5_ R133C with wild type NOTCH3 EGF_1-5_, we obtained excellent in vitro-aggregation,in line with previous reports (Duering *et al*., 2011; Opherk *et al*, 2009), which would increase the chances of producing an immunogenic aggregate. From stable cells lines producing NOTCH3 EGF1-5 from either wildtype (WT) or NOTCH3^R133C^, we purified poly-Histidine- and c-Myc-tagged NOTCH3 EGF_1-5_ WT or NOTCH3^R133C^ fragments (Duering *et al*., 2011) from cell culture medium by metal ion affinity chromatography capturing the poly-Histidine tag (Fig 2A). This resulted in a yield of approximately 90% of the total protein content in the aggregated form (Fig 2B). We next explored the potential of the NOTCH3 EGF_1-5_ peptides to multimerize and produce an amorphous aggregate (Duering *et al*., 2011) by incubating equal amounts of WT with R133C fragments or the WT and R133C fragments separately at 37 °C for 5 days. Spontaneous multimer aggregation increased with time of incubation, which is in line with previous observations (Duering *et al*., 2011; Opherk *et al*., 2009), with a prominent loss of monomeric NOTCH3 EGF_1-5_ when WT and R133C fragments were mixed as compared to incubating them separately (Fig 2C). Aggregates from mixed WT and R133C (WT/R133C aggregates) were therefore selected for the active immunization experiments.

**Figure 2.**
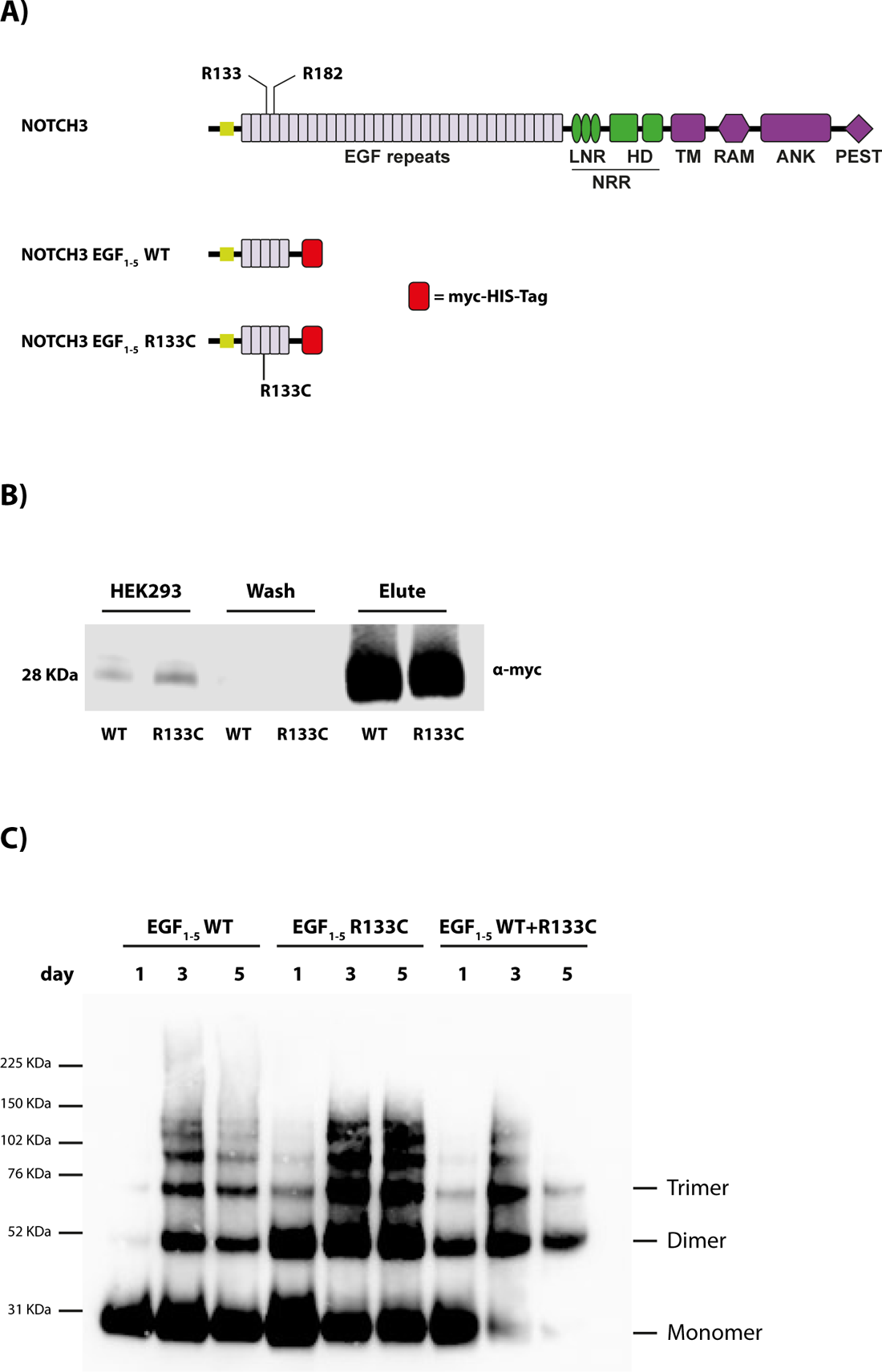
A) Schematic representation of NOTCH3 and NOTCH3 EGF_1-5_. NOTCH3 represents the full-length protein, and NOTCH3 EGF_1-5_ represents the NOTCH3 protein with exon 1 to 5 fused with a myc-His-Tag at the C-terminus used for purification of the aggregated protein. B) Western blot of the NOTCH3 EGF_1-5_ WT and R133C purified protein. The eluate fractions were visualized by western blot using an α-myc antibody. C) Western blot of NOTCH3 EGF_1-5_ WT and R133C aggregated proteins. The incubated fractions of NOTCH3 EGF_1-5_ WT and R133C were visualized on a western blot using an α-myc antibody. The purified proteins and the aggregates were verified after 1-5 days incubation by western blot using α-myc ab under non-reducing conditions.

### N3 EGF_1-5_ antigen evokes a robust immune response in TgN3R182C^150^ mice

We next assessed the immunogenic potential of the aggregated NOTCH3 EGF_1-5_ peptides and whether they could prevent NOTCH3 ECD aggregation. Active vaccination was initiated at three months of age, two months before NOTCH3 protein accumulation is observed in the TgN3R182C^150^ mice (Rutten *et al*., 2015). Aggregated NOTCH3 EGF_1-5_ WT/R133C protein plus adjuvant (with PBS plus adjuvant as sham-immunization control) was used to immunize TgN3R182C^150^ mice at three months of age (Fig 3A) (Kontsekova *et al*, 2014). A booster shot containing aggregated protein plus adjuvant, or PBS plus adjuvant as control, was administered one month later. Two weeks later, another booster shot containing only NOTCH3 EGF_1-5_ WT/R133C protein or PBS was injected, and further booster shots were administered every two weeks until seven months of age, which was the end point of the analysis (Fig 3A) (Kontsekova *et al*., 2014). Using a NOTCH3 EGF_1-5_ aggregate ELISA, we assessed the immune response in the serum of the immunized mice at one, two and four months post-immunization, i.e. at four, five, and seven months of age. We observed a progressive increase in the immune response of the vaccinated versus the sham-immunized mice at one and two months post-vaccination and the increase was further pronounced at four months post-vaccination (Fig 3B). In conclusion, these data show that an immune response is mounted against the injected NOTCH3 EGF_1-5_ WT/R133C aggregates.

**Figure 3.**
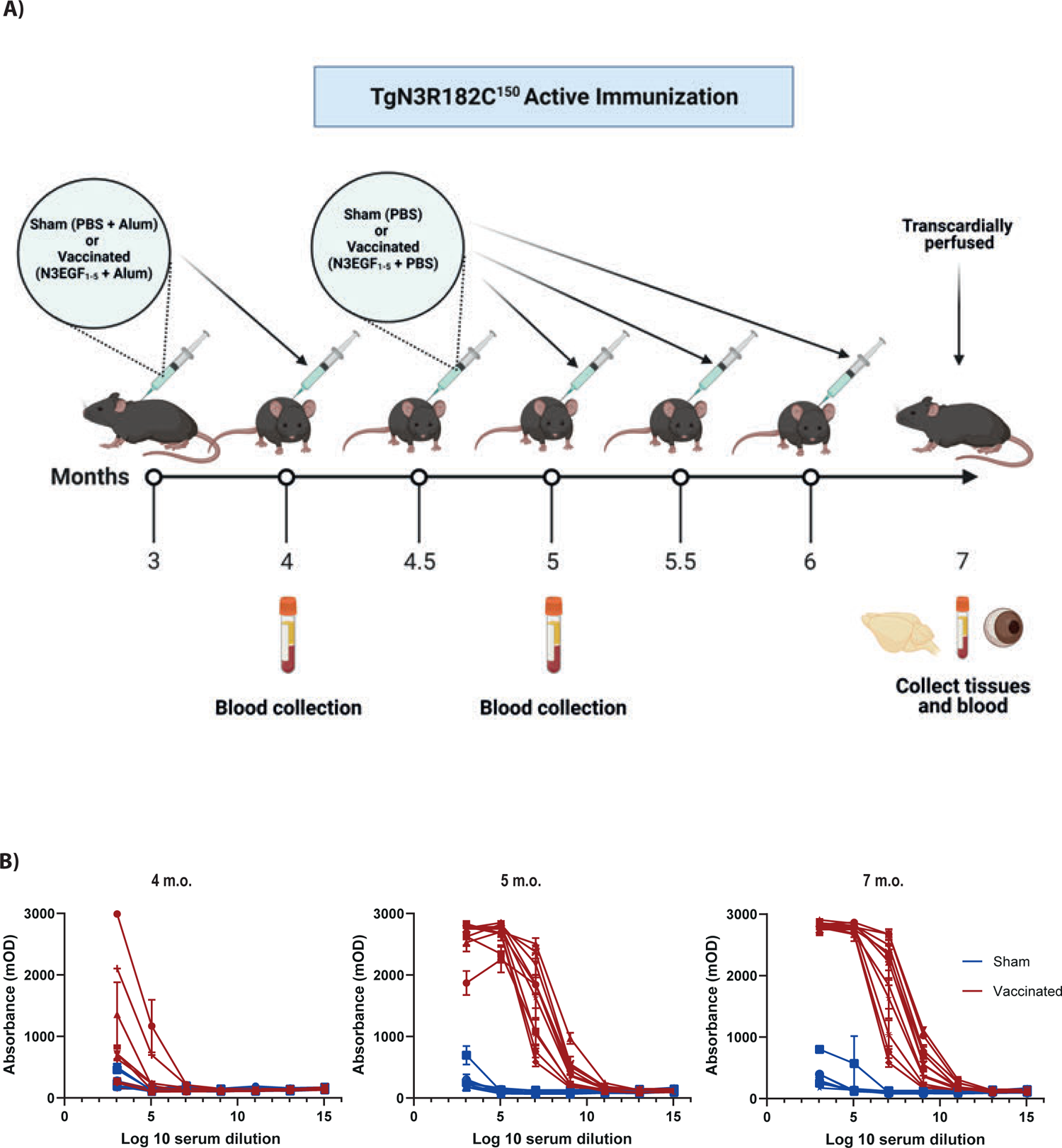
A) Schematic and work plan of the subcutaneous active immunization on the TgN3R182C^150^ mouse model. B) Antibody titre validation of serum from TgN3R182C^150^ CADASIL mice immunized with NOTCH3 EGF_1-5_ aggregates (vaccinated) and PBS (sham) at 4, 5 and 7 months old. A direct ELISA with NOTCH3 aggregate-coated plates and different dilutions of serum was performed.

### The numbers and size of NOTCH3 ECD deposits around capillaries are reduced in NOTCH3 EGF_1-5_ WT/R133C-vaccinated TgN3R182C^150^ mice

We first determined whether active immunization resulted in lower levels of NOTCH3 ECD deposition in the TgN3R182C^150^ brain vasculature. Brain tissue from NOTCH3 EGF_1-5_ WT/R133C-vaccinated TgN3R182C^150^ mice as well as from sham-vaccinated or untreated TgN3R182C^150^ controls was processed for NOTCH3 ECD immunohistochemistry. NOTCH3 ECD deposits have mostly been described to occur in small arteries/arterioles. To assess NOTCH3 accumulation in arteries/arterioles, we stained for NOTCH3 and alpha-smooth muscle actin (ASMA), the latter a marker for arteries/arterioles (Vanlandewijck *et al*, 2018). We observed a significant increase of both the percentage (p = 0.0085) and number (p = 0.0106) of NOTCH3 deposits per vessel area as well as the average size of the deposits (p = 0.0047) in the non-treated 18 months as compared to seven months old TgN3R182C^150^ mice (Fig 4A, B), in keeping with a previous report (Rutten *et al*., 2015). The increase in NOTCH3 deposition was not linked to an increase in expression of the NOTCH3 gene, as no differences in expression of NOTCH3 and three downstream target genes (*Hes1*, *Hey1* and Nrip2) were observed during aging of the TgN3R182C^150^ mice (Fig EV1). We did not observe a difference between the non-vaccinated, sham-vaccinated and vaccinated TgN3R182C^150^ mice at seven months of age. There are several explanations as to why this might be the case. One explanation is that NOTCH3 ECD deposition in arteries/arterioles is not affected by the vaccination. Alternatively, NOTCH3 ECD deposition may start after seven months of age in arteries/arterioles in this mouse model or deposit levels were too low to be successfully abrogated at seven months of age (Fig 4A, B).

**Figure 4.**
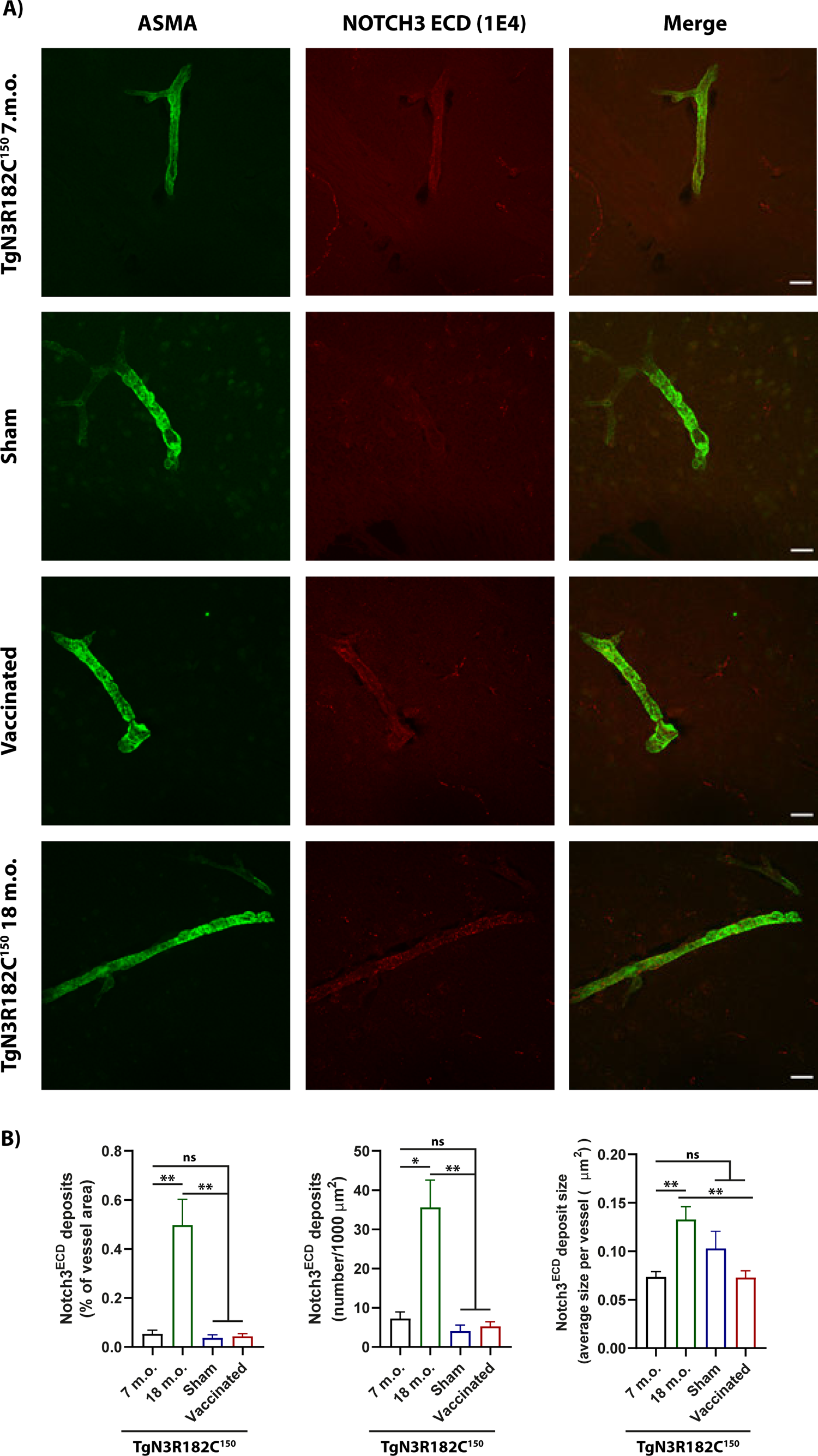
NOTCH3 ECD and smooth muscle actin (SMA) expression in cerebral arteries/arterioles. A) Representative images of TgN3R182C^150^, sham- and NOTCH3 EGF_1-5_ - immunized mice at 7 months of age and TgN3R182C^150^ at 18 months of age. Representative images show brain arteries of TgN3R182C^150^ (7 and 18 months), sham and NOTCH3 EGF_1-5_ - immunized mice stained with a monoclonal antibody against NOTCH3 ECD (1E4, red) and an α-SMA antibody (green). B) Quantification of NOTCH3 ECD deposits (numbers per 1,000 μm^2^) and NOTCH3 ECD stained area and average size per vessel revealed no decrease in NOTCH3 ECD deposition in brain arteries between NOTCH3 EGF_1-5_ - immunized, sham and non-vaccinated TgN3R182C^150^ mice at 7 months of age. NOTCH3 ECD deposits (numbers per 1,000 μm^2^) and NOTCH3 ECD stained area and average size per vessel increases significantly in the TgN3R182C^150^ mice at 18 months of age versus NOTCH3 EGF_1-5_ - immunized, sham and non-vaccinated TgN3R182C^150^ mice at 7 months of age. (*p < 0.05, **p < 0.01, ns= non-significant). Scale bar =20µm.

Given the extensive deposition of NOTCH3 aggregates around capillaries in CADASIL (Yamamoto *et al*, 2013), we next assessed differences in NOTCH3 deposition in the capillaries by using a combination of NOTCH3 and perlecan immunohistochemistry, as perlecan stains the basement membrane around both arteries and capillaries. We found increased NOTCH3 deposition as a percentage of capillary vessel area (p = 0.0032), number of deposits per 1000 µm of capillary vessel area (p = 0.0042), as well as NOTCH3 ECD deposit size (p = 0.0236; Fig 5A, B) in 18-month-old non-treated TgN3R182C^150^ mice compared to their three month counterparts. In contrast to the arteries/arterioles, the increase in the number of NOTCH3 ECD deposits (p = 0.0012) and percentage of deposits per vessel area (p = 0.0001) around capillaries was abrogated in vaccinated TgN3R182C^150^ mice, while no corresponding reductions were noted in the sham-vaccinated TgN3R182C^150^ mice at seven months (Fig 5A, B). Similarly, the average size of NOTCH3 ECD deposits in the vaccinated TgN3R182C^150^ mice was decreased as compared to non-treated or sham-vaccinated mice (p = 0.0047; Fig 5A, B). In conclusion, these data demonstrate that active immunization with NOTCH3 EGF_1-5_ WT/R133C aggregates specifically reduces the amount of NOTCH3 aggregates around cerebral capillaries.

**Figure 5.**
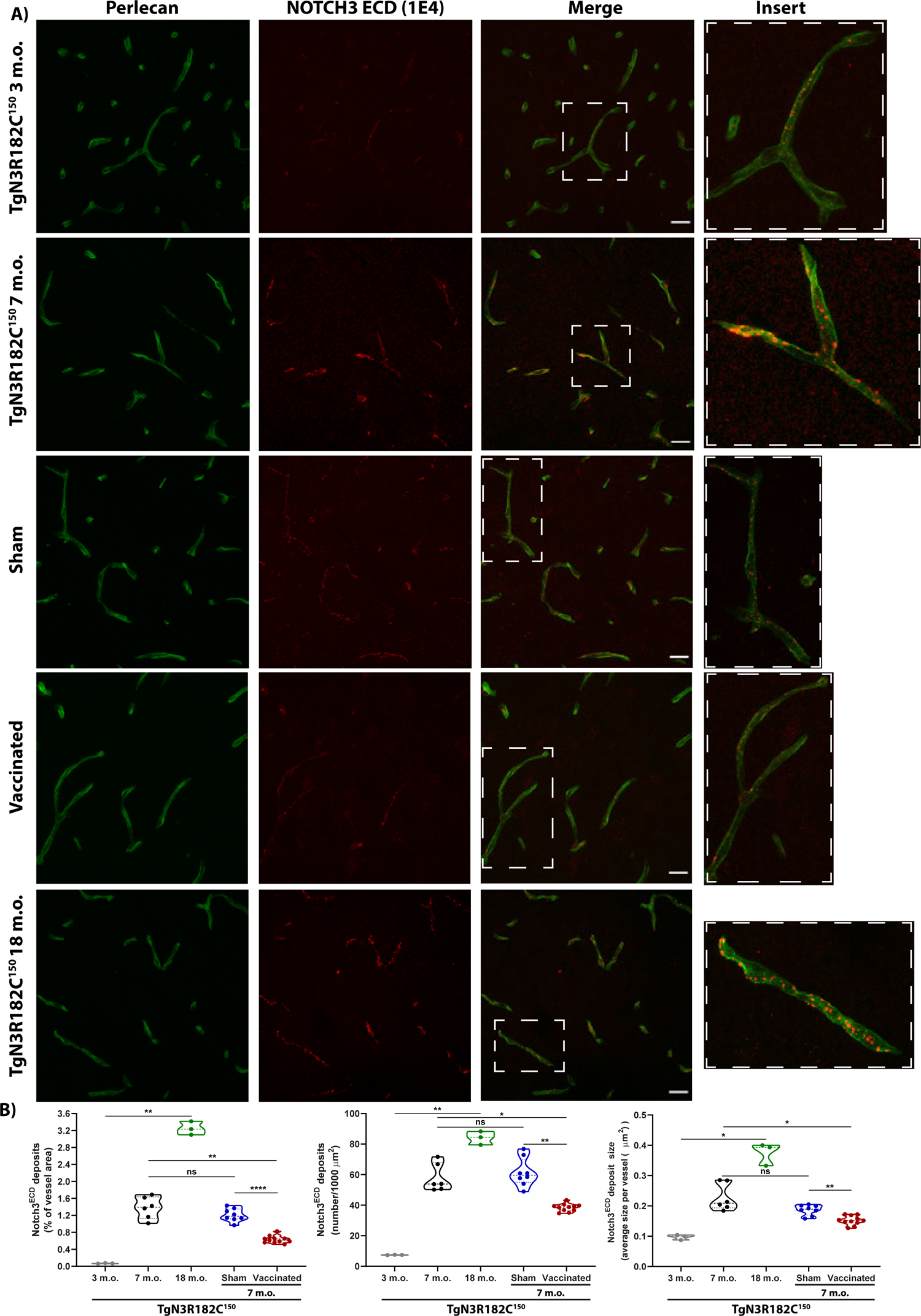
NOTCH3 ECD and perlecan expression in cerebral capillaries. A) Representative images of TgN3R182C^150^, sham- and NOTCH3 EGF_1-5_ - immunized mice at 3, 7 and 18 months of age. Representative images show brain arteries and capillaries of TgN3R182C^150^, sham and NOTCH3 EGF_1-5_ - immunized mice stained with a monoclonal antibody against NOTCH3 ECD (1E4, red) and an anti-perlecan antibody (green). B) Quantification of NOTCH3 ECD deposits (numbers per 1,000 μm^2^) and NOTCH3-ECD stained area and average size per vessel revealed a significant increase in NOTCH3 ECD deposition in brain arteries and capillaries between non-vaccinated 3 months old TgN3R182C^150^ (n=3) and 7 months old TgN3R182C^150^ (n=6) mice and 18 months old TgN3R182C^150^ (n=3). Quantification of NOTCH3-ECD deposits (numbers per 1,000 μm^2^) and NOTCH3-ECD stained area and average size per vessel revealed a significant decrease in NOTCH3-ECD deposition in brain arteries and capillaries between NOTCH3 EGF_1-5_ - immunized (n=11), sham (n=9) and non-vaccinated TgN3R182C^150^ (n=6) mice. (*p < 0.05, **p < 0.01, ****p < 0.0001, ns= non-significant). Scale bar =20µm.

### The amount of NOTCH3 ECD is reduced in blood from NOTCH3 EGF_1-5_ WT/R133C-vaccinated TgN3R182C^150^ mice

Elevated levels of NOTCH3 ECD in the blood have been observed in a CADASIL mouse model and in plasma and serum from CADASIL patients (Primo *et al*, 2016). We therefore wanted to extend the analysis of active immunization to learn whether NOTCH3 ECD blood levels were also altered in the CADASIL mouse model. To assess this, we first measured the levels of NOTCH3 ECD in blood from TgN3R182C^150^ mice using an ELISA assay adapted from a previous report (Primo *et al*., 2016). NOTCH3 ECD was detected in whole blood serum of non-treated TgN3R182C^150^ mice at three months of age and at elevated levels at seven months of age (p = 0.0093; Fig 6A). In contrast, no signal above the background was detected in C57BL/6 WT mice or in Notch3^-/-^ mice at seven months of age (data not shown). Serum NOTCH3 ECD in vaccinated TgN3R182C^150^ mice was significantly reduced compared to sham vaccinated animals (p = 0.0196; Fig 6B). To exclude that the observed reduction of NOTCH3 ECD in the serum was a consequence of competion between the capture antibody in the ELISA and the raised humoral response by the vaccination we added serum from vaccinated C57BL6/J mice to the serum samples from vaccinated TgN3R182C^150^ mice in different dilutions. No difference in the NOTCH3 ECD levels was observed after dilution normalization (data not shown), suggesting that there was no interference between the humoral response and the ELISA capture antibody. Collectively, these data show that active immunization with aggregated NOTCH3 EGF_1-5_ WT/R133C reduces the amount of circulating NOTCH3 ECD in serum in the TgN3R182C^150^ CADASIL mouse model.

**Figure 6.**
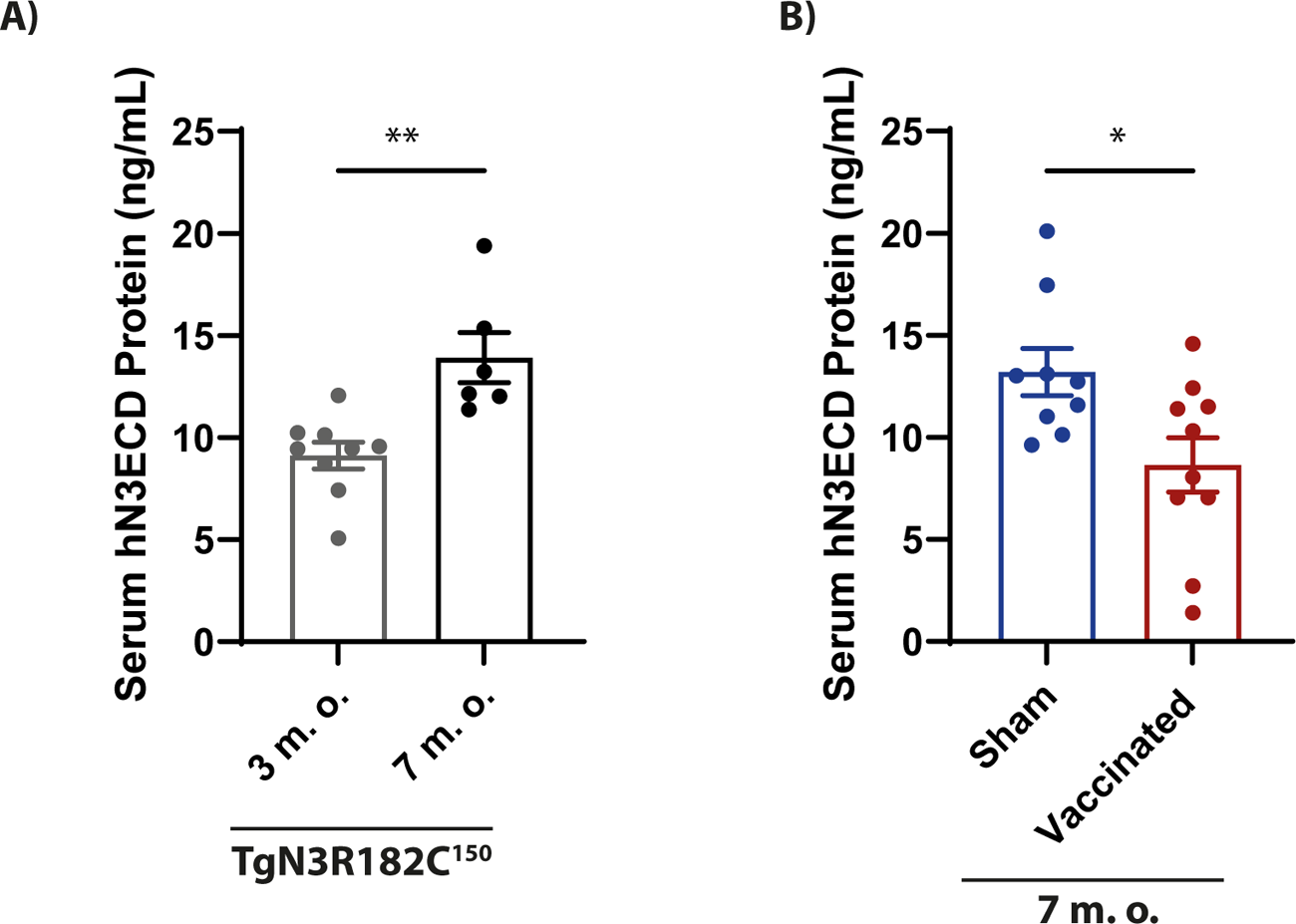
Quantification of human NOTCH3 ECD protein present in the whole blood serum of sham, immunized and non-vaccinated TgN3R182C^150^ mice (at 3 and 7 months old). A) NOTCH3 ECD was detected in the whole blood serum of the non-treated TgN3R182C^150^ mice at three months of age and further increased at seven months of age. B) NOTCH3 ECD in the TgN3R182C^150^ mice was significantly reduced in the vaccinated TgN3R182C^150^ mice. Statistical analysis was performed using unpaired t test with Welch’s correction. P < 0.05 was considered significant (*p < 0.05, **p < 0.01, ***p < 0.001).

### The number of retinal VSMC is not reduced by active immunization

Although the antibodies were generated towards NOTCH3 EGF_1-5_ aggregates, this does not exclude a potential targeting of NOTCH3 monomeric receptors, which may disrupt Notch3 signaling and, in turn, cause adverse effects on the vasculature. The architecture of the retinal blood vessels is particularly amenable to analysis of vascular dysfunction as loss of Notch3 signaling in *Notch3^-/-^* mice leads to a progressive loss of VSMC in the retina (Henshall *et al*, 2015). We therefore wanted to establish whether active immunization with NOTCH3 EGF_1-5_ WT/R133C aggregates reduced the number of VSMC in retinal blood vessels as an indicator of whether Notch3 signalling was disturbed. Immunostaining for smooth muscle actin (ASMA) revealed no significant differences in the smooth muscle cell architecture of the retinal vasculature in control, vaccinated or sham-vaccinated TgN3R182C^150^ mice compared to and WT (C57BL6/J) mice at seven months of age (Fig 7A, B). In contrast, there was extensive loss of VSMC at three months of age in the Notch3^-/-^ mice, compared to three months old WT (C57BL6/J), in keeping with a previous report (Henshall *et al*., 2015) (Fig 7C). Furthermore, the NOTCH3 EGF_1-5_ WT/R133C-vaccinated TgN3R182C^150^ mice were normal with regard to body weight and general behavior (data not shown). Together, this suggests that active immunization against NOTCH3 does not have a detrimental impact on Notch3 signalling.

**Figure 7.**
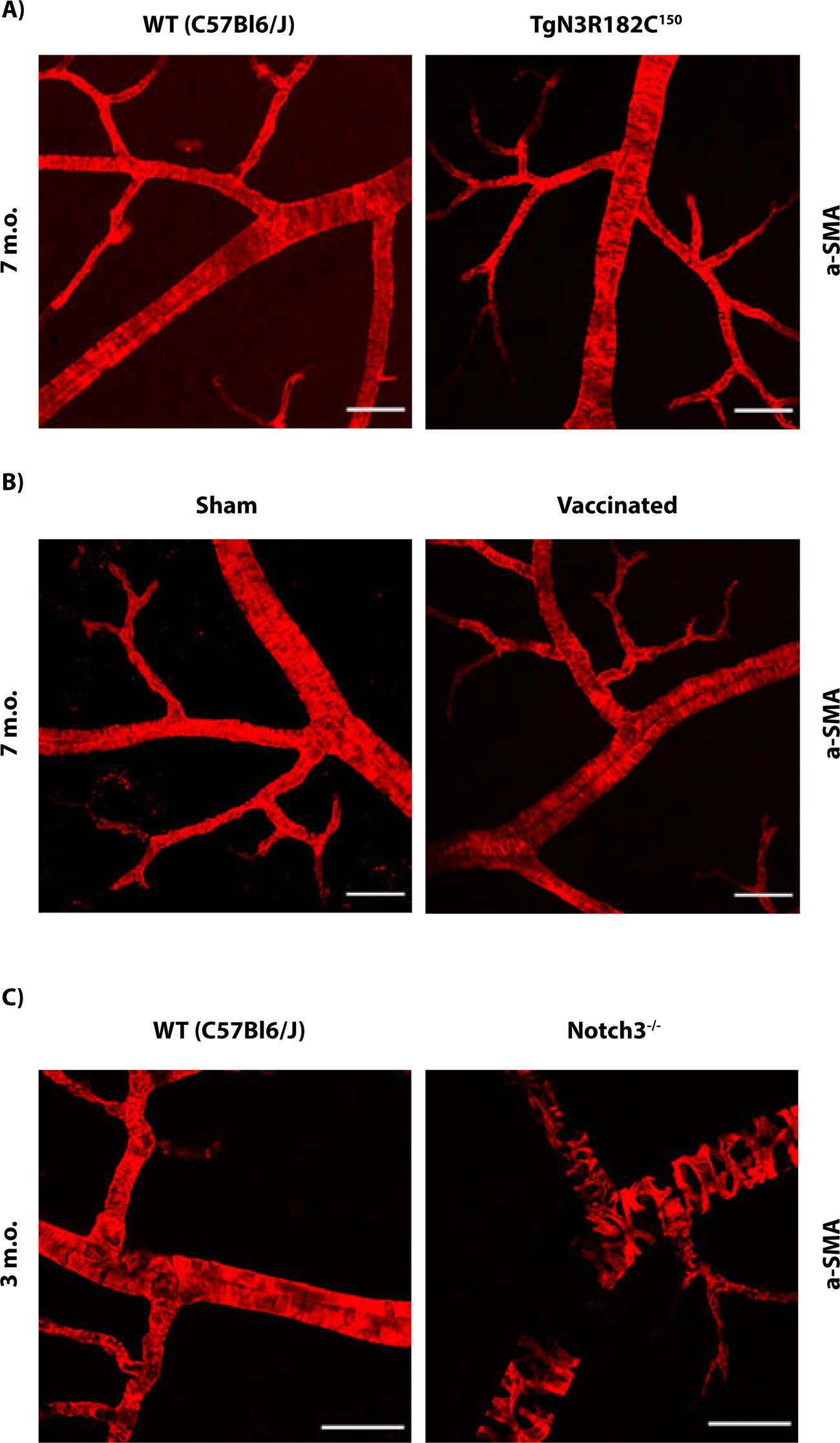
The number of retinal VSMC is not reduced by active immunization. A) Immunostaining for smooth muscle actin (ASMA) revealed that there were no significant differences in the composition of the smooth muscle cell coating of vessels in the retinal vasculature in WT (C57Bl6/J) versus TgN3R182C^150^ mice at 7 months of age. B) Immunostaining for smooth muscle actin (ASMA) shows no significant differences in the composition of the smooth muscle cell coating of vessels in the retinal vasculature in NOTCH3 EGF_1-5_ - vaccinated versus sham-vaccinated TgN3R182C^150^ mice. C) Immunostaining for smooth muscle actin (ASMA) shows an extensive loss of VSMC in the Notch3^-/-^ mice when compared to a WT (C57Bl6/J) at 3 months of age. Scale bar =50µm.

### The number of activated microglia is increased in NOTCH3 EGF_1-5_ WT/R133C-vaccinated TgN3R182C^150^ mice

Microglia are the innate immune cells of the brain and play an important role in phagocytosis of aggregates in the brain (Rogers *et al*, 2002). The reduced deposition of NOTCH3 aggregates in the capillaries of vaccinated TgN3R182C^150^ mice indicates that immune-mediated clearance of these aggregates may been activated, with microglia being a likely culprit. To address this, we performed immunostaining for activated microglia using Iba1 (expressed at low levels in resting microglia and at higher levels in activated and migrating microglia) and CD68 (expressed in activated microglia). Combined immunostaining for Iba1 and CD68 revealed an increase in the number of CD68-positive cells (p = 0.0092) as well as the area (p = 0.0157) in which they were localized in relation to Iba1-staining in seven month old vaccinated TgN3R182C^150^ mice compared to control or sham-vaccinated TgN3R182C^150^ mice (Fig 8A,B). There was also a trend towards more NOTCH3 ECD deposits inside or in close vicinity to microglia in vaccinated TgN3R182C^150^ mice compared to control or sham-vaccinated TgN3R182C^150^ mice, although the difference did not reach statistical significance (Fig 9A, B). In conclusion, the data suggest that active immunization with the NOTCH3 EGF_1-5_ WT/R133C aggregates increases the proportion of microglia with NOTCH3 deposits, while not changing the area or number of deposits within a microglial cell. This may indicate an involvement of microglia in clearing the NOTCH3 ECD aggregates.

**Figure 8.**
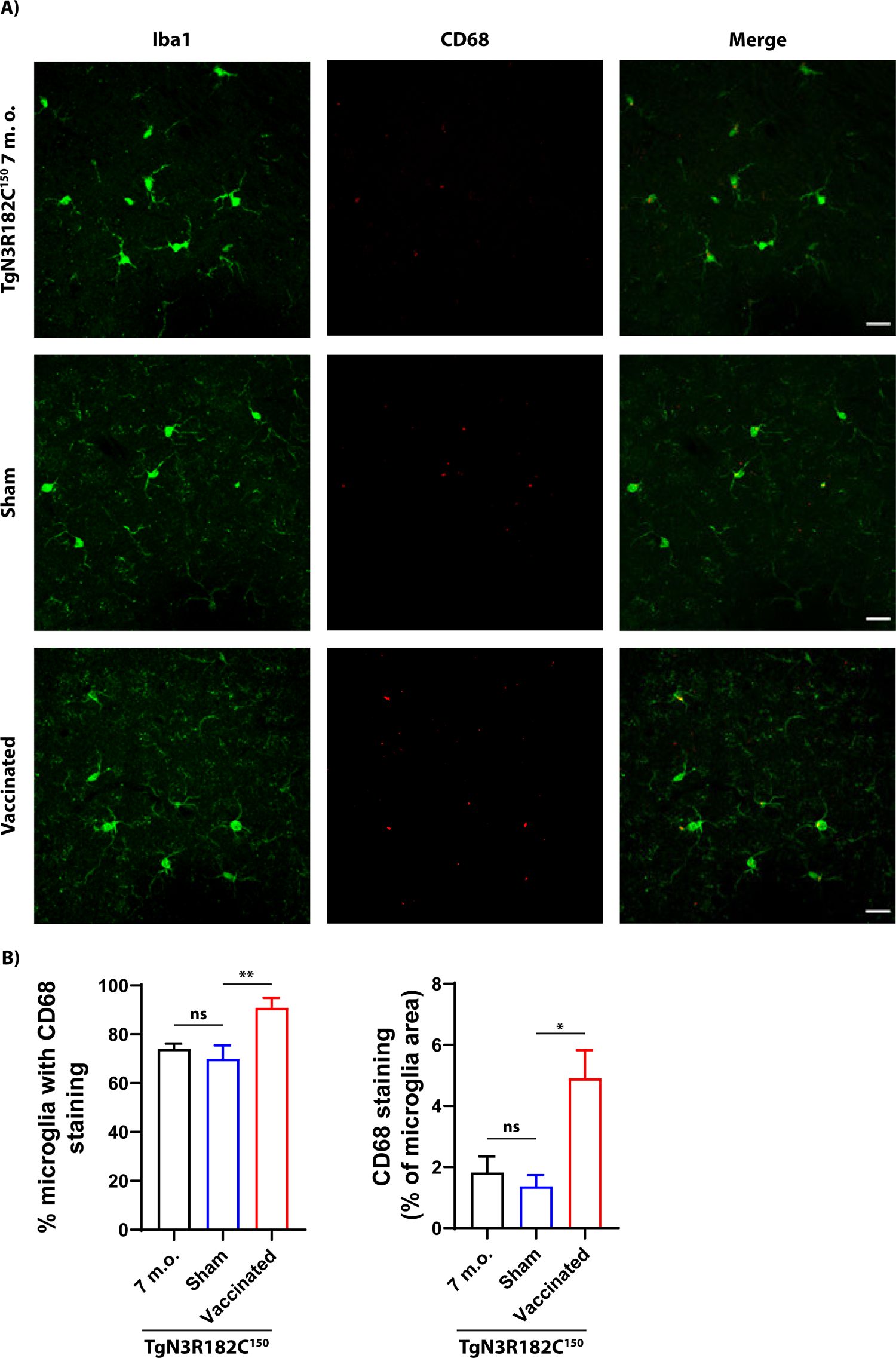
Microglia activation in sham, immunized and non-vaccinated TgN3R182C^150^ mice at 7 months. (A) Representative images show microglia stained with anti-CD68 antibody (red) and Iba1 antibody (green). (B) Quantification of CD68-stained area revealed a significant increase in the % of microglia and microglia area between N3 EGF_1-5_ - immunized (n=11), sham (n=9) and non-vaccinated TgN3R182C^150^ (n=6) mice. (*p < 0.05, **p < 0.01, ns= non-significant). Scale bar =20µm.

**Figure 9.**
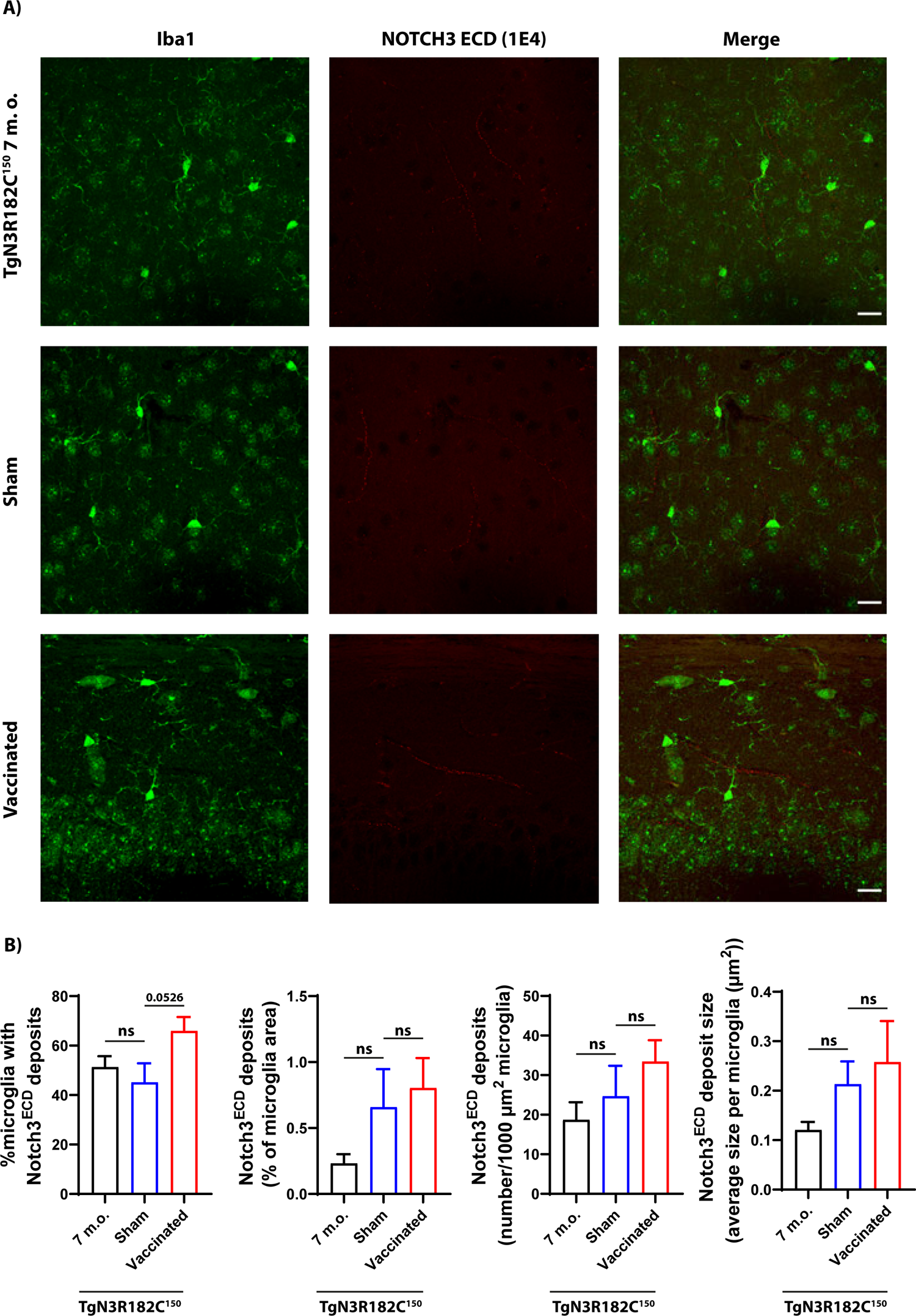
NOTCH3 ECD and microglia expression. A) Representative images of TgN3R182C^150^, sham- and NOTCH3 EGF1-5 - immunized mice at 7 months of age stained with a monoclonal antibody against NOTCH3 ECD (1E4, red) and an antibody against microglia (Iba1, green). B) Quantification of NOTCH3 ECD deposits (numbers per 1,000 μm2) and NOTCH3 ECD stained area and average size per microglia revealed no alterations between the NOTCH3 EGF1-5 - immunized (n=11), sham (n=9) and non-vaccinated TgN3R182C^150^ (n=6) mice at 7 months of age. (ns= non-significant). Scale bar =20µm.

## Discussion

CADASIL is the most common genetic form of small vessel disease but with no available therapies that can prevent, halt or cure the disease. There is thus a need to explore potential therapeutic strategies. In this report we develop a novel active immunization strategy in a preclinical CADASIL model and demonstrate its ability to prevent NOTCH3 ECD aggregation, which is a key hallmark of CADASIL pathophysiology. We find that NOTCH3 ECD depositions around capillaries as well as NOTCH3 ECD levels in the blood are reduced in CADASIL mice treated with the NOTCH3 EGF_1-5_ - directed vaccine for four months and that the vaccine does not appear to interfere with the role of Notch signaling for vascular integrity.

Several lines of evidence indicate that CADASIL pathology stems from aggregation of the NOTCH3 extracellular domain, and likely leads to disruption of the microarchitecture of the cerebrovasculature and unlimately reducing myogenic tone and resulting in a series of lacunar infarcts. Accumulation of NOTCH3 ECD is an early step in the pathogenic process, followed by formation of GOM deposits and the subsequent degeneration of the VSMC (Capone *et al*., 2016; Monet-Lepretre *et al*., 2013). To disrupt NOTCH3 misfolding and aggregation is therefore a promising therapeutic avenue to explore in CADASIL treatment. Immune based therapies have become a leading and prioritized therapeutic strategy for other proteinopathies and neurodegenerative diseases during the last 20 years (Alpaugh & Cicchetti, 2019). In the quest for AD therapies, emerging encouraging results suggest a clinically meaningful effect from immunotherapies aimed at clearing Aβ-amyloid which recently also have led to the first FDA approval of a passive vaccine for treatment of AD (Demattos *et al*, 2012; Golde *et al*, 2009; Knopman *et al*., 2021; Sevigny *et al*, 2016).

Our data provide proof-of-principle that an active immunization strategy can reduce NOTCH3 aggregation in a pre-clinical model of CADASIL and thus may be beneficial also in the treatment of CADASIL. Active immunization against NOTCH3 accumulation using a 1:1 mixture of aggregated recombinantly expressed R133C (475T>C) and wild-type NOTCH3 EGF1-5 truncated protein fragements resuted in significantly reduced deposition of NOTCH3 aggregates in the capillaries of a transgenic CADASIL mouse model. The use of NOTCH3 EGF_1-5_ aggregates containing both mutant and wild-type NOTCH3 as the immunogen was selected as the vast majority of CADASIL patients are heterozygous mutation carriers and thus express both mutated and non-mutated NOTCH3, and it has been demonstrated that misfolded NOTCH3 ECD forms aggregates with wild type NOTCH3 (Duering *et al*., 2011). It is thus reasonable to postulate that wild type NOTCH3 co-aggregates with mutated misfolded NOTCH3 in CADASIL patients, and we noted that there appeared to be a more rapid formation of aggregates when both R133C and wild type NOTCH3 ECD was present.

The reduction in NOTCH3 aggregation contrasts with observation from studies using a passive immunization approach. In one report, use of the mouse monoclonal antibody (5E1) into a mouse model expressing the rat R169C *Notch3* CADASIL mutant revealed robust antibody binding to NOTCH3 aggregates in the brain, improved cerebral blood flow in response to acetylcholine or whisker stimulation, and restoration of myogenic tone of cerebral arteries (Ghezali *et al*., 2018). Interestingly, the amount of NOTCH3 aggregates or GOM was not reduced by 5E1 passive immunization, in contrast to the data presented in our study. Our study aimed to determine whether prevention of NOTCH3 aggregaton in a CADASIL model and did not investigate the capacity of the treatment to resuce vascular pathophysiology mainly due the absence of a CADASIL animal model that fully recapitulates the milieu of symptoms and pathologies seen in CADASIL. Another study was based on the use of an agonistic NOTCH3 antibody (A13) (Li *et al*, 2008) in a mouse model overexpressing C455R *NOTCH3* (Machuca-Parra *et al*., 2017), which generates a hypomorphic form of NOTCH3, leading to loss of VSMC. C455R NOTCH3 mice injected with A13 antibody showed a pronounced recovery of the VSMC coating of retinal vessels at 6 weeks of age, accompanied by a restoration of the decreased levels of NOTCH3 ECD in the blood of the C455R NOTCH3 mice (Machuca-Parra *et al*., 2017). While elegant, an agonist antibody strategy may however only be applicable to CADASIL mutations with hypomorphic signaling, such as C455R, whereas an active immunization strategy may target a broader set of CADASIL mutations, including the aggregation prone R182C, which is signaling neutral (Fig EV2). To facilitate coverage of a broader set of mutations, we opted for using EGF1-5 as an immunogenic agent, as it is the region of NOTCH3 containing most of the CADASIL-causing mutations identified so far (Fig 1B). The fact that our immunization strategy, based on the use of NOTCH3 EGF_1-5_ aggregates containing both wild type and mutated NOTCH3, led to decreased NOTCH3 aggregation around cerebral capillaries and reduced serum levels of NOTCH3 ECD in a mouse model carrying a different *NOTCH3* mutation (R182C) suggests that active immunization with one mutant form can be efficient for CADASIL patients harbouring different types of mutations. Such cross-mutation efficacy is of particular importance for the clinical translatability of a vaccine for CADASIL, as more than 200 cysteine-altering mutations have been identified and thus it would be infeasible to tailor a vaccine for each specific mutation.

The reduction in vascular NOTCH3 deposition was mirrored by reduced serum NOTCH3 ECD levels, providing further support for successful target engagement and suggesting that vascular and serum NOTCH3 levels are linked. This may be in line with a “peripheral sink” model, which posits that there is a passive equilibrium between aggregation-prone NOTCH3 in the blood and the brain, and that “capturing” of the protein in the blood would lead prevent deposition in the brain leading to less aggregation (DeMattos *et al*, 2002; Zhang & Lee, 2011). While theoretically appealing, the peripheral sink hypothesis seems unlikely to operate for Aβ aggregates in Alzheimer’s disease (Georgievska *et al*, 2015; Honig *et al*, 2018), but it will be interesting to learn whether it may be a contender to explain the reduction of Notch3 ECD aggregates following active immunization.

We investigated whether microglia, the resident immune cells of the central nervous system, might be responsible for the reduction in NOTCH3 ECD aggregation in capillaries either by clearing aggregated NOTCH3 or by preventing its aggregation. We did not observe a significant volume of NOTCH3 ECD deposits within microglia, indicating phagocytosis is likely not the main mechanism of NOTCH3 clearance, however this may be a result of rapid turnover in the phagocytosis process. It is also possible that a peripheral humoral immune response leads to increased clearance of NOTCH3 prior to the formation of aggregates in the capillary bed.

As Notch3 signaling plays a pivotal role in vascular smooth muscle cell biology, it is important that NOTCH3-based CADASIL therapies do not affect Notch downstream signaling. In the considerations for the active immunization we reasoned that targeting aggregated NOTCH3 ECD but not monomeric NOTCH3 would be important, and serum from immunized mice indeed showed a strong affinity for aggregated NOTCH3. In support of a preference for aggregated NOTCH3, immunized mice did not display any abnormal behavior or obvious altered phenotype as well as a preserved VSMC coating in the retina, which indicates that endogenous Notch signaling was not affected by vaccination. Furthermore, since MRI is clinically used for CADASIL diagnosis, we have previously analysed a more aggressive CADASIL mouse model (TgN3R182C^350^), which has higher NOTCH3 expression, by MRI and we did not observe any consistent differences in MRI and behaviour between the transgenic model and controls at 20 months of age (Gravesteijn *et al*, 2020; Rutten *et al*., 2015). In the light of these observations, we do not expect the milder CADASIL model used in this study to show any MRI differences at 7 months post-immunization. Together, these data suggest that vaccination with the NOTCH3 EGF_1-5_ vaccine successfully targets NOTCH3 pathology in a tolerable manner.

It will be interesting to further investigate how our active immunization strategy might affect NOTCH3 deposition both long term and in older animals with advanced NOTCH3 aggregation. Our current study was designed to be a proof-of-principle study that demonstrated that active immunization could indeed prevent NOTCH3 deposition. The next steps will be to determine whether it can reverse NOTCH3 deposition and whether being exposed to active vaccination for extended periods of time are safe. It will also be necessary to employ a pre-clinical model of CADASIL that recapitulates the vascular physiology of the disease (i.e. decreased myogenic tone) to determine whether our therapeutic approach extends beyond NOTCH3 aggregate removal. For example, the CADASIL R169C mouse model, which exhibits reduced myogenic tone and cerebral blood flow, would be of interest to further explore our active therapeutic strategy (Ghosh *et al*, 2015).

In conclusion, our study provides proof-of-principle that active immunization is an effective and tolerable therapy to reduce NOTCH3 ECD aggregates around capillaries and reduce NOTCH3 serum levels in a CADASIL mouse model. Given that innocultion with NOTCH3 from one CADASIL variant was capable of reducing aggregate load in another CADASIL variant, this active vaccination therapy could be applicable to a broad spectrum of CADASIL variants. This is the first study demonstrating a reduction of vascular NOTCH3 pathology development and is a promising lead for the development of a CADASIL therapy.

## Material and Methods

### Mouse maintenance, breeding and genetics

TgN3R182C^150^ mice, which overexpress the full length human *NOTCH3* gene from a genomic BAC construct with the archetypal p.Arg182Cys mutation, were generated in a C57BL/6J genetic background. Four mutant strains with different NOTCH3 RNA expression levels relative to endogenous mouse Notch3 RNA expression, including the TgN3R182C^150^ strain, were developed and have been previously described (Rutten *et al*., 2015). Mice were provided with water and food *ad libitum*, were maintained in a 12 hour light/dark cycles, and housed in enriched cages. All experimental animal procedures were performed in accordance with local regulations and rules and were approved by the Stockholm Animal Ethics board (ethical permit No 4433-2020).

### Generation of stable HEK293 NOTCH3 EGF_1-5_ WT and R133C cell lines

We used previously generated constructs encoding a truncated form of human NOTCH3 ECD (WT and R133C) consisting of the first five EGF-like repeats (N3 EGF_1-5_, amino acids 1-234) with a poly-Histidine and c-Myc tags at the C-terminus (Duering *et al*., 2011). From these original constructs we generated codon-optimized constructs using GeneArt gene synthesis (ThermoFisher) in the *piggyBac* transposon system to increase the yield of secreted protein. Human embryonic kidney 293 (HEK293) cells were plated into 6 well plates (0.5×10^6^ cells per well) and cultured in Dulbecco’s modified Eagle’s medium (DMEM; Invitrogen) supplemented with 10% fetal bovine serum (FBS; Invitrogen) and 1% penicillin-streptomycin (Invitrogen) at 37°C in a humidified 5% CO2 atmosphere. The following day 0.5 µg of NOTCH3 EGF_1-5_ WT and R133C plasmids containing the *piggyBac* transposon and 0.2 µg of *piggyBac* transposase were transfected using Lipofectamine 2000 (Invitrogen) according to the manufacturer’s instructions. 24 hours after transfection, cells were split and selected for two weeks in DMEM medium containing 1 mg/mL of G418 (Invitrogen). After two weeks, the selected cells were expanded and maintained in DMEM medium with 0.6 mg/mL of G418.

### Purification of NOTCH3 EGF_1-5_ WT and R133C proteins

We purified NOTCH3 EGF1-5 WT and R133C proteins as previously described (Oliveira *et al*, 2022). Briefly, HEK293 NOTCH3 EGF_1-5_ WT and R133C (N3 EGF_1-5_ WT and R133C) cells were grown in DMEM with 10% FBS until near confluency and were then washed with Dulbecco’s phosphate-buffered saline (DPBS) and supplemented with DMEM medium without FBS for 5 days. The conditioned medium from both cell lines was collected, cleared from cell debris by centrifugation at 1500 g and dialyzed with the aid of SnakeSkin Dialysis Tubing 10 KDa MWCO (ThermoFisher) in PBS for 24 hours at 4 °C. Dialysed media from 3 flasks (∼150 mL) of each cell line was equilibrated with 1 ml (bead volume) of cobalt ion affinity resin (TALON Superflow, GElifesciences) for 30 minutes with agitation at 4 °C. After the resin solution was added on a gravity flow column and washed three times with wash buffer (200 mM Sodium Phosphate buffer, 500 mM NaCl and 5 mM imidazole, at pH 7.5). After elution with elution buffer (200 mM Sodium Phosphate buffer, 500 mM NaCl and 300 mM imidazole, at pH 7.5) the fractions were pooled and dialyzed in a dialysis membrane (Spectrum™ Spectra/Por™ 1 6-8 KDa MWCO, FisherScientific) against PBS for 24 h at 4 °C. The pooled fractions of the NOTCH3 EGF1-5 WT and R133C were concentrated to 1 mg/mL in a concentration column (Vivaspin 6 10 KDa MWCO, Sartorius) according to the manufacturer’s instructions. The protein concentration was calculated by measuring the concentrated proteins with a BCA absorbance assay according to the manufacturer’s instructions (Pierce™ BCA Protein Assay Kit #23225, ThermoFisher).

### Immunization of TgN3R182C^150^ mice with aggregated NOTCH3 EGF_1-5_ protein

NOTCH3 EGF_1-5_ WT and NOTCH3 EGF_1-5_ R133C proteins were incubated 1:1 to a final concentration of 0.5 mg/mL in PBS at 37^0^C for 5 days with gentle agitation (350 rpm), and the presence of aggregated NOTCH3 EGF_1-5_ protein was verified using western blot and SDS-PAGE under non-reducing conditions. The aggregated NOTCH3 EGF_1-5_ protein was mixed with the adjuvant or PBS to a concentration of 0.25 mg/mL. We performed the active immunization using a similar immunization schedule as Kontsekova and colleagues (Kontsekova *et al*., 2014) with slight modifications adjusted to using mouse rather than rat and the size of the immunized protein Briefly, 20 three months old transgenic TgN3R182C^150^ mice were immunized initially either with 25 µg of aggregated NOTCH3 EGF_1-5_ protein (n=11, vaccinated) plus adjuvant (Imject Alum Adjuvant, ThermoFisher) or PBS (n=9, sham) plus adjuvant, followed by a booster shot four weeks later containing 25 µg of aggregated NOTCH3 EGF_1-5_ protein plus adjuvant and the corresponding sham booster shot. Subsequent booster shots of sham and aggregated NOTCH3 EGF_1-5_ protein (25 µg) in PBS were performed every two weeks until a total of six immunization events have taken place over a period of four months.

### Whole blood serum collection

Whole blood serum was collected from the immunized mice (n=20), and non-immunized TgN3R182C^150^ at 3 months (n=9) and 7 months (n=6). Briefly, mice were anesthetized intraperitoneally with avertin (240 mg/kg) and blood was then collected from the right ventricle and dispensed to a tube with serum gel with clotting activator (Microvette 500 Z-Gel, Sarstedt). After centrifugation for 5 min at 10000x g, the whole blood serum was collected, aliquoted and stored at −80 °C.

### Antibody immune response titer validation of whole blood serum by ELISA

To detect the immune response of the immunization, we designed an indirect enzyme-linked immunosorbent assay (ELISA) against the NOTCH3 EGF_1-5_ aggregates. Briefly, the plates were coated with the antigen (NOTCH3 EGF_1-5_ aggregates) at 0.625 µg/mL, followed by incubation overnight at 4 °C. Blocking solution was added (PBS-T with 1% BSA and 10% goat serum) for 2 hours at room temperature (RT). Next, the plates were washed 2 times with PBS, and the diluted whole blood serum samples of both sham and vaccinated mice were added to the plates followed by incubation overnight at 4 °C. The following day, the plates were washed 4 times with PBS and incubated with polyclonal anti-mouse HRP conjugated secondary antibody (1:2000) for 2 hours at RT. After washing five times with PBS, the plates were incubated for 30 minutes with Pierce 1-Step Ultra TMB ELISA Substrate (ThermoFisher). Pre warmed sulfuric acid stop solution (DY994, R&D Systems) at 37 °C was used to stop the reaction and the absorbance was monitored at 450 nm with a Fluorostar galaxy plate reader.

### NOTCH3 ECD custom ELISA

To detect NOTCH3 ECD in circulation, we designed an NOTCH3 ECD sandwich ELISA based on a previously described protocol (Primo *et al*., 2016). Briefly, high-affinity binding 96 well plates were coated with a NOTCH3 capture monoclonal antibody (MAB1559; R&D Systems) at 0.625 ng/µL in 100 µL of PBS and agitated overnight at 4 °C. The plates were blocked for 3 hours with agitation in a 10% BSA solution at RT. Whole blood serum samples were diluted 1:40 in 100 µL of reagent diluent (DY995; R&D Systems) and recombinant human NOTCH3 ECD (1559-NT-050; R&D Systems) was used as standard protein. The samples and standard were added to the plates and incubated with agitation overnight at 4 °C. After three washes with washing buffer (WA126, R&D systems), 100 µL of a solution containing 0.001 mg/mL of the detection biotinylated polyclonal antibody (BAF1559; R&D Systems) raised against NOTCH3 ECD and 2.5% BSA was added and was used for detection followed by agitation for 2h at RT. The plates were washed 3 times with washing buffer and a horseradish peroxidase-streptavidin (HRP-Strep) complex (R&D Systems DY998) diluted 1:25 in reagent diluent was added to the plates and incubated for 40 minutes with agitation. After incubation and washing 5 times, the plates were incubated for 30 minutes with Pierce 1-Step Ultra TMB ELISA Substrate (ThermoFisher). Pre-warmed sulfuric acid stop solution (DY994, R&D Systems) at 37 °C was used to stop the reaction and absorbance at 450 nm was measured in a Fluorostar galaxy plate reader.

### Brain and retina collection and histology

Mice were deeply anesthetized intraperitoneally with avertin (240 mg/kg) and transcardially perfused with 50 mL of DPBS followed by 50 mL of 4% paraformaldehyde (PFA). The brain and retinas were collected and were postfixed overnight in 4% PFA at 4°C. One hemisphere of the brain was paraffin-embedded and was sagittal serial sectioned in 5-10 µm sections and placed in positively charged slides.

### Immunohistochemistry

Sections were deparaffinized in xylene 2 times during 5 min, followed by rehydration of the tissue through a series of graded alcohols (100% ethanol (2x for 3 min), 95% ethanol (1x for 3 min), 70% ethanol (1x for 3 min) and H_2_O MQ (1x 3 min)). Next, heat antigen retrieval was performed by submerging the slides in Diva Decloaker buffer (Biocare Medical) and placing them in a pressure cooker (Biocare Medical) for 30 minutes at 110 °C. Slides were cooled at RT for 30 min and placed in PBS for 15 min on a shaker. Brain sections were outlined with a hydrophobic barrier using a PAP pen, and a permeabilization solution (PBS with 2% BSA and 0.3% Triton-X) was loaded onto the tissue for 10 min at RT. The tissues were then blocked for 30 min with Rodent Block M (Biocare Medical) followed by 30 min with blocking solution (PBS with 1% BSA and 10% goat serum). To assess the NOTCH3 ECD deposits the tissues were incubated overnight at 4 °C with mouse monoclonal anti-human NOTCH3 ECD primary antibody (1:100, clone 1E4, Millipore) followed by detection with Alexa 594 conjugated anti-mouse secondary antibody (1:1000, Life Technologies). Capillaries were identified by immunostaining with rat monoclonal anti perlecan antibody (1:100, clone A7L6, Millipore) overnight at 4 °C followed by detection with Alexa 488 conjugated anti-rat secondary antibody (1:1000, Life Technologies). Arteries were identified by immunostaining with a mouse monoclonal anti actin, α-Smooth Muscle - FITC antibody (1:500, clone 1A4; Sigma-Aldrich) overnight at 4 °C. Microglia was stained with rat monoclonal anti-CD68 antibody (1:100, Bio-Rad) and with a rabbit monoclonal anti-Iba1 antibody (1:100, WAKO) overnight at 4 °C followed by detection with Alexa 488 conjugated anti-rabbit secondary antibody (1:1000, Life Technologies), and Alexa 594 conjugated anti-rat secondary antibody (1:1000, Life Technologies). Stained sections were imaged at 63x magnification using a Zeiss LSM880 microscope. Images were captured with identical settings across sections. The entire procedure was performed under blinded conditions.

### Quantification of NOTCH3 ECD deposits

The quantitative image analyses were performed blinded to the genotype and using pre-established parameters (ImageJ software, Fiji distribution) (Schindelin *et al*, 2012). We analyzed NOTCH3 ECD deposits on maximal intensity projections of image stacks by applying the following pipeline: (1) manual delineation of arteries and delineation of capillaries by automated segmentation on the perlecan channel, and subsequent creation of a region of interest (ROI) for each vessel followed by the measurement of the vessels area; (2) on the NOTCH3 ECD channel the segmentation was achieved by performing automatic image thresholding (RenyiEntropy method), followed by automatic detection and counting of number and the area of NOTCH3 ECD deposits inside each ROI by using the “Analyze Particles” function of Fiji. To quantify the NOTCH3 ECD deposits three nonadjacent sections (50 µm apart) were analyzed per mouse (10-14 images randomly selected per animal section). The mean of the randomly selected image fields (224.92 × 224.92 μm) per section was used to quantify NOTCH3 ECD load in capillaries and arteries. All vessels (capillaries and arteries) in the randomly selected image fields (224.92 × 224.92 μm) were included in the quantification of NOTCH3 ECD deposits as an ROI. Results are represented as the number, average size and surface area of NOTCH3 ECD deposits on the vessel area.

### Quantification of activated microglia

The quantitative image analysis was performed blinded to the genotype as described above. We analyzed the activated microglia on maximal intensity projections of image stacks by applying the following pipeline: (1) delineation of microglia by automated and manual segmentation on the Iba1 channel, and subsequent creation of a ROI for each microglia followed by the measurement of the microglia area; (2) on the CD68 channel (activated microglia) the segmentation was achieved by performing automatic image thresholding (RenyiEntropy method), followed by automatic detection and counting of number and the area of CD68 staining inside each ROI by using the “Analyze Particles” function of Fiji. To quantify the activated microglia three nonadjacent sections (50 µm apart) were analyzed per mouse (4-6 images randomly selected per animal section). The mean of the randomly selected image fields (224.92 × 224.92 μm) per section was used to quantify CD68 stain in the microglia area. All microglia in the randomly selected image fields (224.92 × 224.92 μm) were included in the quantification of activated microglia as an ROI. The % of microglia area represents the percentage that the CD68 staining (stained by Iba1) occupies per microglia area. To analyze the NOTCH3 ECD deposits that colocalize with the microglia area, we employed the same pipeline described above to select the microglia area in the Iba1 channel. We then measured the NOTCH3 ECD deposits in the ROI employing the same pipeline we used for measuring the load of NOTCH3 ECD deposits in the vessels. All microglia in the randomly selected image fields (224.92 × 224.92 μm) were included in the quantification of NOTCH3 ECD deposits as an ROI. Results are represented as the number, average size and surface area of NOTCH3 ECD deposits on the microglia area.

### Immunostaining of whole mount retina

The mouse retinas were dissected and removed from the eyecups and washed three times in PBS at RT for five minutes, followed by permeabilization in 0.3% TritonX-100 in PBS and blocked in PBS with 2% BSA and 10% goat serum for 2 hours at RT. Retinas were then incubated for 24 hours at 4°C with primary antibodies diluted in PBS containing 3% BSA, 0.1% Triton and with secondary antibodies diluted in the same buffer. After washes with 0.1% Triton X-100/PBS, retinas were flat mounted and imaged using a Zeiss LSM880 microscope.

### Luciferase assay

NIH3T3 cells were transfected with 12×CSL-luc and CMV-β-galactosidase together with *NOTCH3* wildtype, R182C plasmid or pcDNA3 vector as a control. Five hours post transfection, cells were seeded on ligand Jag2 or Fc coated plates and DMSO as a vehicle and the γ-secretase inhibitor DAPT (10 µM) were added overnight. After incubation for 24 hours, the cells were lysed in Cell culture lysis reagent (Promega) and luciferase activity was measured in triplicate by the GloMax® Multi Detection System apparatus (Promega) using the Dual-Glo Luciferase Assay System (Promega) according to the manufacturer’s protocol. In all assays, relative luciferase activity was calculated as the ratio of Luciferase values normalized to β-gal levels.

### Quantitative real-time PCR analysis

Brain vessels were isolated as previously decribed (Matthes *et al*, 2021), followed by RNA extraction using RNeasy mini kit (QIAGEN) and cDNA synthesis with Maxima First Strand cDNA Synthesis Kit (ThermoFisher Scientific). Real-time PCR analysis was performed using the Applied Biosystems 7500 Fast Real-Time PCR System according to the manufacturer’s protocol. The primers for qPCR are Hes1: forward 5’-CCAGCCAGTGTCAACACGA-3’, reverse 5’-AATGCCGGGAGCTATCTTTCT-3’; Hey1: forward 5’-CCCCTCACCCTACTCACCA-3’, reverse 5’-GCTTCAACCCAGACCCAA-3’; Nrip2: forward 5’-GGGAAGAATGTGTGGAGTGG-3’, reverse 5’-GAAGGCAGCAATGAAGAAGC-3’; NOTCH3: forward 5’-GAAGTTACCCCCAAGAGGCAA, reverse 5’-TATCTCGGTCACGCTGCAA-3’.

## Statistical analysis

GraphPad Prism 9 was used to generate the graphs, All results are presented as mean ± standard error of the mean (SEM). Shapiro-Wilk normality and lognormality and F tests were performed before choosing the statistical test to apply. Student’s t-test was used to assess the statistical differences between experimental groups. Multiple comparisons were evaluated by 1-way analysis of variance (ANOVA) followed by Brown-Forsythe and Welch ANOVA tests and followed by Dunnett’s T3 multiple comparisons test in cases where different groups did not have equal variance. P < 0.05 was considered significant (*p < 0.05, **p < 0.01, ***p < 0.001, ****p < 0.0001).

---

**Figure EV1.**
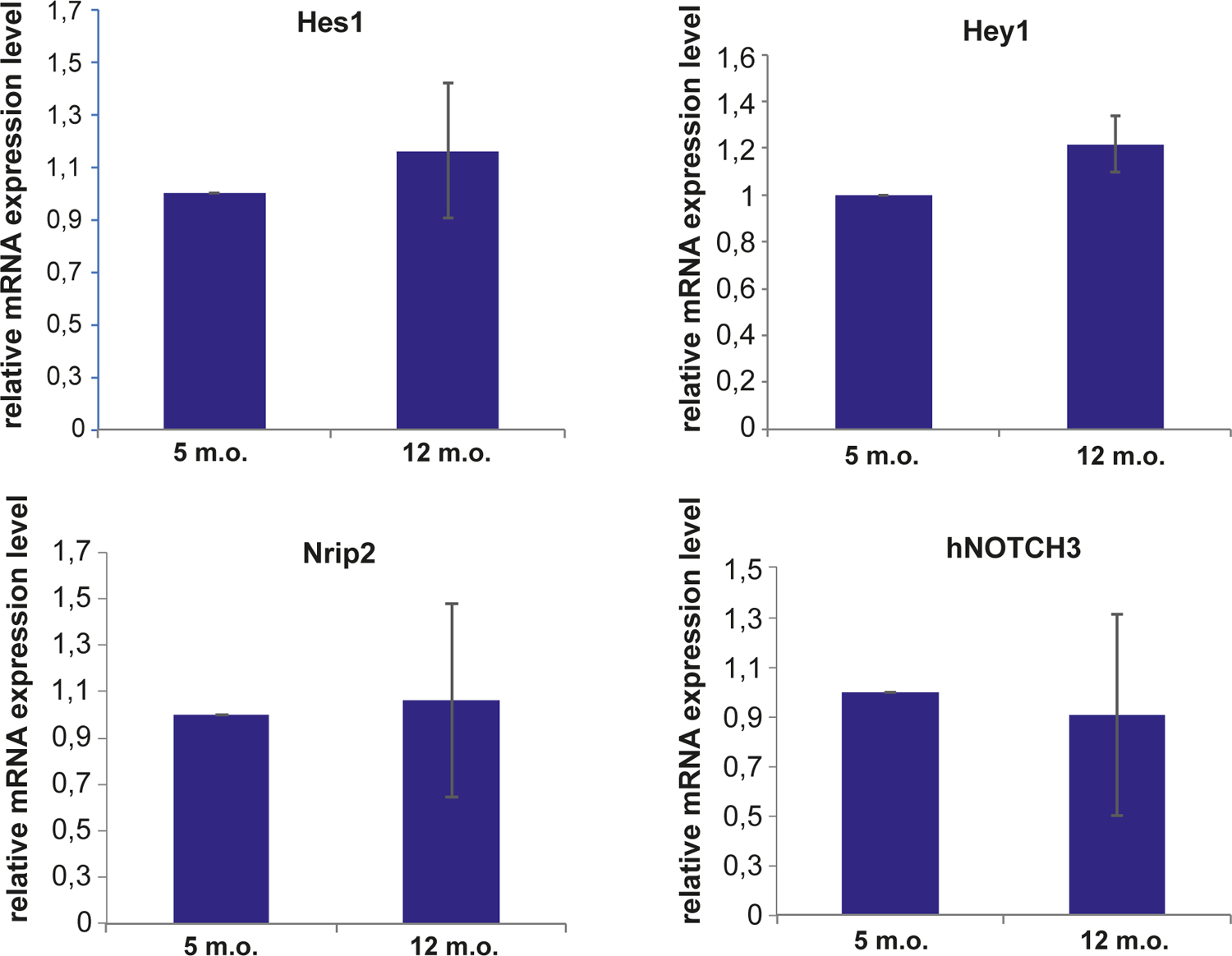
Quantitative real-time PCR analysis of the Notch downstream target genes *NOTCH3, Hes1, Hey1* and Nrip2 on TgN3R182C150 mice at 5 and 12 months of age.

**Figure EV2.**
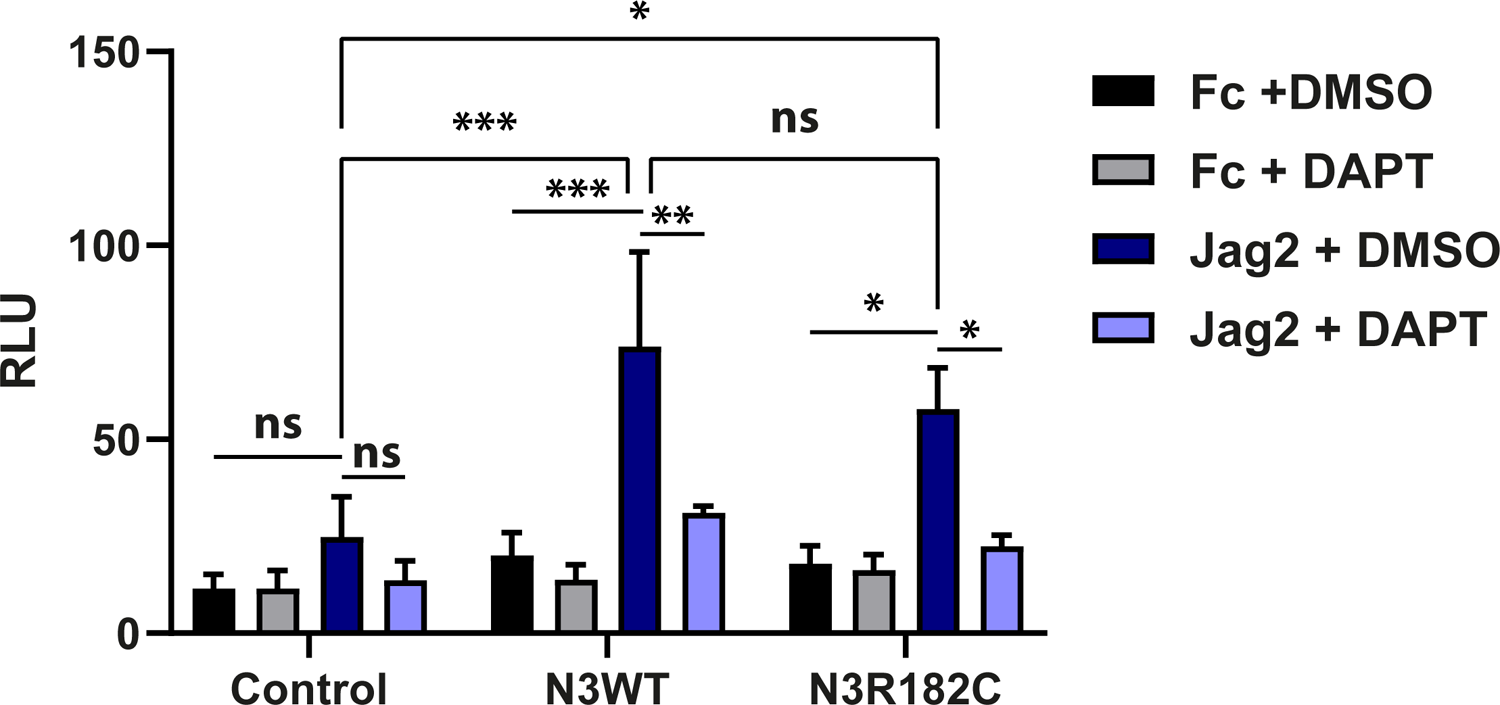
NIH3T3 cells were transfected with the control, wild type NOTCH3, or NOTCH3 R182C plasmids, as well as the β-gal and 12XCSL-luc reporter plasmids and cultured on immobilized jagged2 (Jag2) in the presence of DMSO or DAPT (n=3 and two technical replicates). Statistical analysis was performed using 2-way ANOVA followed by Tukey’s multiple comparisons tests. P < 0.05 was considered significant (*p < 0.05, **p < 0.01, ***p < 0.001, ns= non-significant). RLU, relative luminescence units.

## Acknowledgments

The authors thank Gido Gravesteijn for help in the custom NOTCH3 ECD ELISA. Some of the figures were created with biorender.com. The financial support from HjärtLungfonden, Hjärnfonden, Gamla Tjänarinnor stiftelse, Gun och Bertil Stohnes stiftelse, Postdoc JUNIOR Fund of Charles University, Swedish Alzheimer Foundation, StratNeuro KI and the Swedish Research Council (2019-00285) is gratefully acknowledged.

